# In vitro assessment of the capacity of pesticides to act as agonists/antagonists of the thyroid hormone nuclear receptors

**DOI:** 10.1101/2020.12.21.423731

**Authors:** Yanis Zekri, Laure Dall Agnol, Frédéric Flamant, Romain Guyot

**Affiliations:** Romain Guyot Institut de Génomique Fonctionnelle de Lyon, Univ Lyon, CNRS UMR 5242, INRAE USC 1370 École Normale Supérieure de Lyon, Université Claude Bernard Lyon 1, 46 allee d’Italie F-69364 Lyon, France

## Abstract

Several in vitro tests, including transcriptome analysis of neural cells, were performed to assess the capacity of 33 pesticides to act as thyroid hormone disruptors (THD). Although some pesticides elicit a cellular response, which interferes with thyroid hormone signaling, we found no evidence that they can act as receptor agonists or antagonists. We conclude that the nuclear receptors of thyroid hormone are not common targets of THD, and that pesticide neurodevelopmental toxicity is not explained by a general alteration of neural cell response to thyroid hormone.

## Introduction

It is now recognized that a number of environmental chemicals act as endocrine disruptors. Among these, thyroid hormone disruptors (THD) interfere with the thyroid system, and have broad adverse effects. In particular neurodevelopment is highly sensitive to a deficit (Gilbert *et al.* 2020) or an excess (Laureano-Melo *et al.* 2019) of thyroid hormone (TH). Therefore early life exposure to THD might have irreversible consequences on cognitive functions. A number of non-exclusive mechanisms have been proposed to explain the adverse effects of THD, which can be classified in two categories:

- Some chemicals interfere with the synthesis or degradation of TH. A number of underlying mechanisms have been documented: xenobiotics can alter the hypothalamus-pituitary-thyroid axis regulation, inhibit iodine uptake, impair the TH production of the thyroid gland or accelerate the catabolism of TH. The outcome of all these processes is a reduction in the circulating levels of TH or an unbalance between T4 (thyroxine) and T3 (3,3’,5’-Triiodothyronine), the most active form of TH. In exposed rodents, this is often associated to histological alterations of the thyroid gland.
- Other chemicals might interfere with the intracellular pathway by which T3 activates the transcription of genes in many cell types. T3 mainly acts in cell nuclei by binding to the C-terminal domain of nuclear receptors (TRs, including TRα1, TRβ1, and TRβ2). Upon T3 binding, the conformation of the C-terminal domain changes (Yen *et al.* 2006). As a result, chromatin-bound TRs release transcription corepressors, and recruit transcription coactivators, up-regulating the transcription of a number of genes (Perissi *et al.* 2005). Like other endocrine disruptors (Toporova *et al.* 2020), THD might compete with T3 to bind the C-terminal domain of TRs and exert an agonist or antagonist influence on T3 signaling (Gorini *et al.* 2020, Guyot *et al.* 2014, Ibhazehiebo *et al.* 2010, Ibhazehiebo *et al.* 2011, Freitas *et al.* 2011). Although such an interference with the cellular response to T3 is expected to have major adverse effect, it does not necessarily modify the circulating levels of TH. TR agonists or antagonists might thus impair neurodevelopment at doses where they do not necessarily modify the circulating levels of TH.

The risk associated to the second class of THD, which only alter the cellular response to T3, is thus better assessed in cultured cells than in rodents, where toxicological tests mainly consider the modes of action from the first class.

Epidemiological associations and toxicological studies indicate that a number of pesticides have a detrimental influence on neurodevelopment. For this reason, they are suspected to act as THD (Leemans *et al.* 2019). As TH circulating levels or thyroid gland histology are rarely altered (Paul *et al.* 2012, Moser *et al.* 2015, Yaglova *et al.* 2017) while direct binding to TRs can sometimes be observed at high concentration (Xiang *et al.* 2017) it is tempting to speculate that pesticides belong to the second class of chemicals, which interfere with the cellular response to T3. If this hypothesis is correct, the danger of pesticides exposure would be widely underestimated under the current registration procedures.

We performed here an *in vitro* assessment of the capacity of common pesticides to act as TRα1 agonists or antagonists. We used a combination of three cell assays to limit the risk of false positives. Combined with genome-wide analyses of gene expression, our data indicate that, unlike estrogenic disruptors, THD do not frequently act as agonists or antagonists of the nuclear receptors of TH, but sometimes exert a partial and indirect influence on T3 response.

## Material and methods

### Chemicals, cell culture medium and toxicity assays

All chemical solutions were prepared by dissolving purified compounds (Sigma Aldrich St Louis MI USA) in dimethylsulfoxide (DMSO). Cells were cultivated in GlutaMAX Dulbecco modified Eagle medium (GlutaMAX DMEM, Thermo Fisher Scientific) with 10% (HEK293) or 12% (C17.2 or SHSY5Y) of newborn calf serum (Thermo Fisher Scientific). Endogenous TH were depleted from the serum by stripping with activated charcoal (Sigma Aldrich, St Louis MI USA) to prevent background activation of reporter constructs. The toxicity of each chemicals was tested on each cell line using the CellTiter-Glo Luminescent Cell Viability Assay (Promega, Madison WI, USA).

### Transactivation in neural cells

The C17.2α-Hrluc reporter cell line was described previously (Guyot *et al.* 2014). It was derived from murine neural stem cells transfected to overexpress in a stable manner the mouse TRα1 receptor. A reporter construct was also introduced in the genome, in which the gene encoding *Renilla luciferase* is driven by 5426 nt of the promoter region of the *Hr/Hairless* gene, which is highly sensitive to T3 transactivation (Thompson 1996). T3 and/or tested compounds were added in the medium 24 hours after seeding cells in 24 well-plates (10^5^ cells/well). DMSO was used in control cells to keep solvent concentration constant. Cells were lysed after 24 h of chemical exposure, and luciferase activity was measured in cell lysates (Renilla luciferase assay system, Promega Madison WI, USA).

### One-hybrid transactivation assay

The HEK293–Gal4TRα1luc cell line was described previously (Guyot *et al.* 2014). It is derived from human HEK293 cells and integrates two DNA constructs. One is encoding a hybrid receptor in which the DNA binding domain of the yeast Gal4 transcription factor is fused to the ligand binding domain of mouse TRα1 receptor. The second carries the firefly luciferase reading frame, driven by a Gal4 responsive promoter with 5 UAS elements. HEK293–Gal4TRα1luc cells were seeded in 24 well-plates (10^5^ cells/well). T3 and tested compounds were added in the medium 24 h later. DMSO was used in control cells to keep solvent concentration constant. Cells were lysed 48h after seeding and luciferase activity was measured with the firefly Renilla luciferase assay system (Promega Madison WI, USA).

### Two-hybrid corepressor interaction

The assay was performed in HEK293 cells transfected for the transient expression of several constructs, as described before (le Maire *et al.* 2020). The pBKGal4NcoR construct encodes a Gal4NcoR hybrid protein, which normally acts as a transcription repressor on expression vectors driven by the UAS DNA binding elements. The pBKVP16TRα1 construct encodes the VP16 transactivation domain of a trans-acting protein from an herpes simplex virus fused to the ligand-binding domain of mouse TRα1. pGal4REx5-βglob-luc-SVNeo construct is an UAS driven luciferase reporter. In this setting, the interaction between Gal4NcoR and the unliganded VP16TRα1 hybrid protein results in an activation of luciferase expression. Addition of T3 destabilizes the interaction, resulting in a reduction in luciferase activity. The pBK-βgal plasmid was also included, which drives the permanent expression of the E coli lacZ. This enabled to use β-galactosidase activity as an internal standard to correct for any variation in transfection efficiency. HEK293 cells were seeded in 24 well-plates (10^5^ cells/well). Cells were transfected the following day with 100ng of DNA containing a mixture of the 4 plasmids (20ng Gal4REx5-βglob-luc-SVNeo, 30ng pBKGal4NcoR, 30ng pBKVP16TRα1, 20ng pBK-βgal) (Markossian *et al.* 2018, Gurnell *et al.* 2000) with the TransIT-Lt1 transfection reagent (Mirus Corporation Madison WI, USA). T3 and tested compounds were added in the medium 4-6h later. DMSO was used in control cells to keep solvent concentration constant. Cells were lysed 24 h after chemical exposure to measure luciferase activity (Firefly luciferase assay system; Promega Madison WI, USA) and β-galactosidase activity, using ortho-nitrophényl-β-galactoside as substrate (β-Galactosidase Reporter Gene Activity Detection Kit, Sigma St Louis MI USA).

### qRT-PCR analysis

C17.2α neural cells, expressing TRα1 (Chatonnet *et al.* 2013), were seeded in 6-wells plates (3.10^5^ cells/well). T3 and the tested compounds were added in the medium the next day. DMSO was used in control cells to keep solvent concentration constant. Cells were lysed 24 h after chemical exposure and RNA extracted using a Macherey-Nagel NucleoSpin RNA II kit. RNA concentrations were measured with a Nanodrop spectrophotometer (Thermo Fisher Scientific). Each RNA sample was reverse transcribed using murine leukemia virus reverse transcriptase (Promega) and random DNA hexamer primers. Quantitative PCR was performed according to a standard protocol, using the Biorad iQ SYBRGreen kit and the CFX96 thermocycler (Biorad). Hprt, a housekeeping gene, was used as an internal control. For each pair of primers (5’CAGCGTCGTGATTAGCGATG + 5’CGAGCAAGTCTTTCAGTCCTGTCC for *Hprt*, 5’CAGCGTCGTGATTAGCGATG + 5’AGAGGTCCAAGGAGCATCAAGG for *Hr*, 5’CACGCCTCCGAAAAGAGGCACAA + 5’ CTTTTCCCCAGTGTGGGTCCGGTA for *Klf9*) a standard curve was established and PCR efficiency was controlled to be within usable range (90%–110%) before analysis using the 2^−ΔΔ(Ct)^ method (Livak *et al.* 2001).

### Transcriptome analysis

RNA were extracted from either human SHSY5Y (ATCC^®^ CRL-2266™) cells or mouse C17.2α (Chatonnet *et al.* 2013) exposed to chemical and/or T3 (10^−9^M or 10^−8^M). Two methods were used for transcriptome analysis: Ion AmpliSeq™ Transcriptome Gene Expression Kit (Thermo Fischer Scientific) run on an Ion Proton™ Sequencer, and RNAseq. In the later case, cDNA libraries were prepared using the total RNA SENSE kit (Lexogen, Vienna Austria) and deep sequencing was performed on a NextSeq500 (Illumina) sequencer as described (Guyot *et al.* 2014, Richard *et al.* 2020). Count tables were prepared using htseq-count (Galaxy Version 0.6.1galaxy3) (Anders *et al.* 2015). Differential gene expression analysis was performed with DEseq2 (Galaxy Version 2.1.8.3) (Love *et al.* 2014) using two factors (pesticide and T3 treatments) and the following thresholds: p-adjusted value < 0.05; expression > 10 reads per million. Clustvis (https://biit.cs.ut.ee/clustvis/) was used for clustering analysis, using the Ward unsquared method and Euclidian distances to prepare heatmap representations.

### Gene Set Enrichment Analysis

Gene Set Enrichment Analysis (Mootha *et al.* 2003, Subramanian *et al.* 2005) was performed with the GSEA Software (version 4.1.0) of University of California San Diego (https://www.gsea-msigdb.org/gsea/index.jsp) using default parameters. We calculated for each compound an Enrichment Score (ES) representing the overrepresentation of genes in the ranked list of up- or down-regulated genes. We retained as significant the compounds inducing a coordinated disruption of thyroid hormone response genes expression with a nominal pval ≤ 0.05.

## Results

### Transactivation assays in neural cells

A set of 33 pesticides was selected among pesticides, which are of common or limited use in agriculture (Medina-Pastor *et al.* 2020, Crepet *et al.* 2013) and covers the main class of pesticides: carbamates, neonicotinoids, organochlorines, organophosphates, pyrethroids, quinolones, strobilurins and triazols. Two reporter cell lines were used to test their ability to interfere with T3 signaling (Guyot *et al.* 2014) with complementary advantages: C17.2α-Hrluc cells are murine cells of neural origin. They host a reporter construct based on the natural promoter of the *Hairless* gene, a well-characterized target gene of TRα1. HEK293–Gal4TRα1luc cells derive from a human fetal kidney cell line. They express a hybrid receptor, which can upregulate an UAS-luciferase reporter, also integrated in the cell genome, after binding of T3 to the C-terminal domain of TRα1. Although this artificial setting does not take into account all the molecular events, which can potentially interfere with TRα1-mediated transactivation, it is also more sensitive to T3. We first determined the maximal tolerable concentration for the two reporter cell lines, defined as 10% of the lowest concentration with visible toxicity. Two inhibitory TRα1 ligands were used as positive controls (Suppl. figure S1): NH-3 prevents the interaction of TRα1 with both coactivators and corepressors (Nguyen *et al.* 2002) while 1-850 is a competitive antagonist (Schapira *et al.* 2000). Each reporter cell line was used in two modes: either in the absence or presence of T3, to respectively identify TRα1 agonists and antagonists. In each case, negative controls, which were performed with equal amount of solvent were used as reference. Overall the variability of the results was ±10% which allowed to define thresholds beyond which luciferase activity was considered as significantly altered. A subset of the selected pesticides was active on C17.2α-Hrluc (Table 1) and or HEK293–Gal4TRα1luc cells (Table 2) either in the absence of the presence of T3. In these assays, both 1-850 and NH-3 act as antagonists, reducing luciferase activity in the presence of T3 and acting in a dose dependent manner. By contrast, active pesticides exert a negative influence on luciferase activity in both the absence and presence of T3. This suggests that these are not true TRα1 antagonists but unspecific inhibitors of the reporter systems, which should be considered as false positives.

**Table 1:** Antagonist activity of selected chemicals on C17.2α-Hrluc reporter cells.

| Molarity of Chemical | 10-5M |  | 10-6M |  | 10-7M |  | 10-8M |  |
| --- | --- | --- | --- | --- | --- | --- | --- | --- |
| T3 10-8M | - | + | - | + | - | + | - | + |
| 1-850 | **0.76±0.05** | **0.59±0.05** | 1.03±0.11 | 0.90±0.04 | 1.05±0.07 | 0.90±0.06 |  |  |
| NH3 | **0.70±0.08** | **0.39±0.04** | 1.04±0.07 | **0.70±0.04** | 1.06±0.03 | 0.81±0.06 | 1.08±0.05 | 0.92±0.04 |
| Azoxystrobin | **0.40±0.02** | 0.85±0.07 | 0.99±0.2 | 1.04±0.26 | 1.24±0.13 | 1.07±0.12 |  |  |
| Benoxacor | 1.09±0.16 | 1.01±0.08 | 0.95±0.15 | 0.95±0.12 |  |  |  |  |
| Bifenthrin | **0.59±0.10** | **0.82±0.10** | **0.69±0.10** | 1.15±0.12 | 0.84±0.04 | 1.10±0.07 |  |  |
| Bitertanol | 0.87±0.09 | 1.07±0.02 | 0.86±0.03 | 1.02±0.17 | 0.88±0.01 | 1.20±0.11 | 0.89±0.06 | 0.87±0.09 |
| Captafol |  |  | 0.95±0.16 | 1.00±0.16 | 0.89±0.07 | 0.97±0.17 |  |  |
| Captan |  |  | 1.01±0.06 | 0.79±0.13 | 1.02±0.13 | 0.93±0.12 |  |  |
| Chlorothalonil |  |  | **1.44±0.05** | 0.84±0.06 | 1.15±0.2 | 0.93±0.05 |  |  |
| Chlorpyrifos | **0.66±0.21** | **0.63±0.07** |  |  |  |  |  |  |
| Chlorpyrifos methyl | **0.67±0.15** | **0.64±0.17** |  |  |  |  |  |  |
| Cypermethrin | 0.81±0.07 | **0.65±0.13** |  |  |  |  |  |  |
| Dichlorodiphenyltrichloroethane |  |  | 1.06±0.05 | 0.92±0.06 |  |  |  |  |
| Deltamethrin | **0.78±0.04** | **0.78±0.21** |  |  |  |  |  |  |
| Dieldrin |  |  | **0.60±0.11** | **0.71±0.16** | **0.83±0.03** | 0.96±0.17 | 0.83±0.08 | 1.10±0.22 |
| Dienochlor | 0.88±0.14 | **0.72±0.04** | 0.95±0.17 | 0.99±0.06 | 1.04±0.13 | 1.02±0.01 |  |  |
| Dinoseb |  |  | **0.75±0.10** | 0.79±0.14 |  |  |  |  |
| Emamectin Benzoate |  |  | **0.79±0.06** | **0.67±0.08** | 0.94±0.1 | 0.83±0.04 |  |  |
| Fenithroton | **0.53±0.04** | **0.49±0.06** | 0.89±0.10 | 0.90±0.07 | 0.91±0.08 | 1.06±0.21 |  |  |
| Folpet | 1.09±0.17 | 0.98±0.21 | 1.09±0.09 | **0.79±0.07** |  |  |  |  |
| Imidacloprid | **0.67±0.09** | **0.72±0.09** | **0.77±0.06** | 0.84±0.12 | **0.70±0.12** | 0.90±0.33 |  |  |
| Malathion | **0.73±0.13** | **0.71±0.08** |  |  |  |  |  |  |
| Penconazol | **0.48±0.04** | **0.23±0.001** | 0.95±0.10 | 1.04±0.07 | 1.03±0.09 | 1.04±0.07 |  |  |
| Phosalone * | 1.00±0.05 | 0.55±0.07 | 1.17±0.08 | 1.16±0.03 | 1.27±0.10 | 1.09±0.03 |  |  |
| Picoxystrobin |  |  | **0.61±0.1** | **0.49±0.01** | 1.19±0.11 | **0.57±0.24** |  |  |
| Piperonyl Butoxide | **0.46±0.07** | **0.49±0.03** | **0.45±0.05** | **0.57±0.01** | **0.69±0.08** | **0.69±0.15** |  |  |
| Prothioconazol |  |  | 1.25±0.12 | 1.13±0.02 | 1.21±0.15 | 1.18±0.1 | 1.15±0.19 | 1.30±0.13 |
| Pyraclostrobin |  |  | **0.53±0.01** | 0.88±0.21 | **0.79±0.06** | **0.64±0.1** | 0.94±0.03 | 0.93±0.15 |
| Pyridaben |  |  | **0.45±0.01** | **0.45±0.11** |  |  |  |  |
| Quinoxifen | **0.44±0.04** | **0.44±0.10** | **0.55±0.01** | **0.60±0.02** | **0.72±0.05** | **0.68±0.05** |  |  |
| Tau-fluvalinate | 1.09±0.26 | **0.76±0.03** |  |  |  |  |  |  |
| Tris(1,3-dichloro-2-propyl)phosphate |  |  | 1.28±0.16 | 0.98±0.04 | 1.28±0.21 | 1.14±0.02 | 1.26±0.19 | 1.16±0.04 |
| Thiram |  |  | 1.29±0.29 | 1.19±0.06 |  |  |  |  |
| Triclosan * | 0.94±0.07 | **0.74±0.06** | 1.11±0.07 | 0.83±0.08 | 1.00±0.12 | 0.83±0.02 | 1.06±0.12 | 0.83±0.04 |
| Trifloxystrobin |  |  | **0.64±0.07** | **0.66±0.04** |  |  |  |  |
| Ziram |  |  | 1.14±0.26 | 1.05±0.10 |  |  |  |  |

Data are the relative activity compared to a matched control with equal concentration of the DMSO solvent (0.1%) \* Actual concentration is 2-fold inferior to the one indicated. Bold characters indicate deviation from controls, by at least 10%.

**Table 2:** Antagonist activity of selected chemicals on HEK293–Gal4TRα1luc reporter cells.

| Molarity of Chemical | 10-5M |  | 10-6M |  | 10-7M |  | 10-8M |  |
| --- | --- | --- | --- | --- | --- | --- | --- | --- |
| T3 10-9M | - | + | - | + | - | + | - | + |
| 1-850 | **0.70±0.02** | **0.23±0.02** | 0.89±0.02 | 0.97±0.07 | 0.91±0.01 | 1.02±0.05 | 0.93±0.06 | 0.96±0.00 |
| NH3 |  |  | **1.44±0.2** | **0.53±0.07** | 1.03±0.03 | 0.90±0.03 | 0.94±0.06 | 0.94±0.01 |
| Azoxystrobin | **0.69±0.02** | **0.51±0.02** | 1.12±0.24 | 1.09±0.12 | 1.04±0.16 | 0.89±0.07 | 1.15±0.09 | 0.80±0.03 |
| Benoxacor |  |  | 1.07±0.08 | 0.99±0.03 | 0.95±0.09 | 0.95±0.03 | 0.79±0.13 | 0.92±0.06 |
| Bifenthrin | 0.96±0.13 | 1.12±0.02 |  |  |  |  |  |  |
| Bitertanol |  |  | 0.89±0.08 | 0.97±0.04 | 0.87±0.04 | 0.99±0.05 | 0.87±0.03 | 0.94±0.05 |
| Captafol |  |  | 0.98±0.02 | 0.84±0.14 | 0.80±0.1 | 0.66±0.04 | 0.63±0.15 | 0.70±0.05 |
| Captan |  |  | 0.96±0.04 | 0.84±0.07 | 0.90±0.01 | 0.70±0.02 | 0.86±0.06 | 0.84±0.04 |
| Chlorothalonil |  |  | **0.35±0.05** | **0.31±0.06** | **0.38±0.06** | **0.33±0.03** | 0.71±0.08 | 0.34±0.08 |
| Chlorpyrifos | **0.89±0.01** | 0.96±0.11 |  |  |  |  |  |  |
| Chlorpyrifos methyl | 0.95±0.13 | 0.88±0.02 |  |  |  |  |  |  |
| Cypermethrin | 1.01±0.01 | 0.89±0.13 |  |  |  |  |  |  |
| Dichlorodiphenyltrichloroethane |  |  |  |  | 0.88±0.08 | 1.11±0.12 |  |  |
| Deltamethrin | **0.80±0.02** | **0.80±0.07** |  |  |  |  |  |  |
| Dieldrin* | 1.05±0.07 | 1.38±0.10 | 1.16±0.04 | 1.31±0.07 |  |  |  |  |
| Dienochlor |  |  | 0.89±0.01 | **0.55±0.07** |  |  |  |  |
| Dinoseb |  |  | **0.81±0.08** | **0.87±0.09** | **0.76±0.09** | **0.78±0.02** | 1.29±0.03 | 1.05±0.12 |
| Emamectin Benzoate |  |  | 0.92±0.09 | **0.82±0.05** | 0.93±0.07 | **0.76±0.10** | 0.92±0.10 | 0.89±0.09 |
| Fenithiotron* | 1.04±0.20 | 1.16±0.17 | 1.03±0.06 | 0.99±0.12 |  |  |  |  |
| Folpet |  |  | 0.98±0.04 | 0.91±0.07 | 0.95±0.07 | 0.83±0.08 | 0.87±0.04 | 0.82±0.05 |
| Imidacloprid | 1.01±0.16 | 1.04±0.02 |  |  |  |  |  |  |
| Malathion | 0.96±0.06 | 0.97±0.01 |  |  |  |  |  |  |
| Penconazol* | 1.08±0.16 | 0.99±0.04 | 0.87±0.09 | 0.99±0.09 |  |  |  |  |
| Phosalone* | **0.68±0.10** | **0.65±0.02** | 1.04±0.1 | 1.14±0.07 |  |  |  |  |
| Picoxystrobin |  |  | **0.81±0.05** | 0.94±0.10 | 0.95±0.27 | 1.09±0.04 | 1.02±0.05 | 1.25±0.29 |
| Piperonyl Butoxide | 1.27±0.02 | 1.04±0.21 |  |  |  |  |  |  |
| Prothioconazol |  |  | 0.88±0.04 | 1.21±0.15 | 0.87±0.10 | 1.15±0.03 | 0.93±0.08 | 1.16±0.06 |
| Pyraclostrobin |  |  | 1.01±0.11 | **0.54±0.01** | 0.98±0.07 | 0.97±0.14 | 0.98±0.11 | 0.99±0.15 |
| Pyridaben | 0.86±0.06 | **0.80±0.01** | **0.82±0.02** | **0.86±0.05** | **0.82±0.04** | **0.76±0.02** | 0.90±0.08 | 0.95±0.01 |
| Quinoxifen * | 1.20±0.06 | **0.65±0.05** | 0.96±0.09 | 0.91±0.40 |  |  |  |  |
| Tau-fluvalinate | 1.07±0.04 | 1.11±0.09 |  |  |  |  |  |  |
| Tris(1,3-dichloro-2-propyl)phosphate |  |  | **0.68±0.10** | 0.97±0.29 | **0.67±0.05** | 0.95±0.06 | **0.70±0.02** | 0.80±0.16 |
| Thiram |  |  |  |  | 0.95±0.10 | **0.45±0.06** | 0.92±0.06 | **0.60±0.21** |
| Triclosan | 1.03±0.04 | 1.09±0.10 | 1.02±0.06 | 1.08±0.08 | 0.96±0.14 | 1.07±0.08 | 0.98±0.10 | 1.03±0.04 |
| Trifloxystrobin |  |  | 0.92±0.04 | 1.15±0.11 | 0.83±0.06 | 1.13±0.07 | 1.02±0.12 | 0.84±0.07 |
| Ziram |  |  |  |  | **0.49±0.02** | **0.52±0.02** | 1.01±0.09 | 0.82±0.02 |

Data are the relative activity compared to a matched control with equal concentration of the DMSO solvent (0.1%) \* Actual concentration is 2-fold inferior to the one indicated. Bold characters indicate deviation from controls, by at least 10%.

### Two-hybrid corepressor interaction

We reasoned that the existence of such false positives would be identified with a third assay in which the addition of antagonists increases, rather than decreases, the luciferase activity. This was achieved in a two-hybrid assay, in which cells were transfected to test the interaction between two hybrid proteins: the Gal4NcoR, which binds DNA at UAS elements, and the VP16TRα1 transactivator. In this setting, the addition of T3 results in the destabilization of the interaction between the two proteins. By contrast to what happens in test 1 and 2, T3 produces a reduction of luciferase activity (le Maire *et al.* 2020, Bochukova *et al.* 2012). Reciprocally, supplementation with the TR antagonist 1-850 increases the luciferase activity in presence of T3 (Suppl. figure S1). We ran this assay for the pesticides that were active in the previous assays. None of the tested product behaved as a TRα1 antagonist in this assay (Table 3). However some of the chemical were active in this test. Interestingly, the influence of piperonyl butoxide seemed to be influenced by the presence of T3. Pyraclostrobin activity also showed some similarity to the one of NH-3. We also used RT-qPCR to address more directly the influence of pesticides on C17.2α cells, at the mRNA levels of 2 well-characterized TRα1 target genes: *Klf9* and *Hairless* (Table 4). While we expected that *Hairless* mRNA level would mirror the results of the first test (Table 1) this was only the case for the reference compounds, 1-850 and NH-3.

**Table 3:**
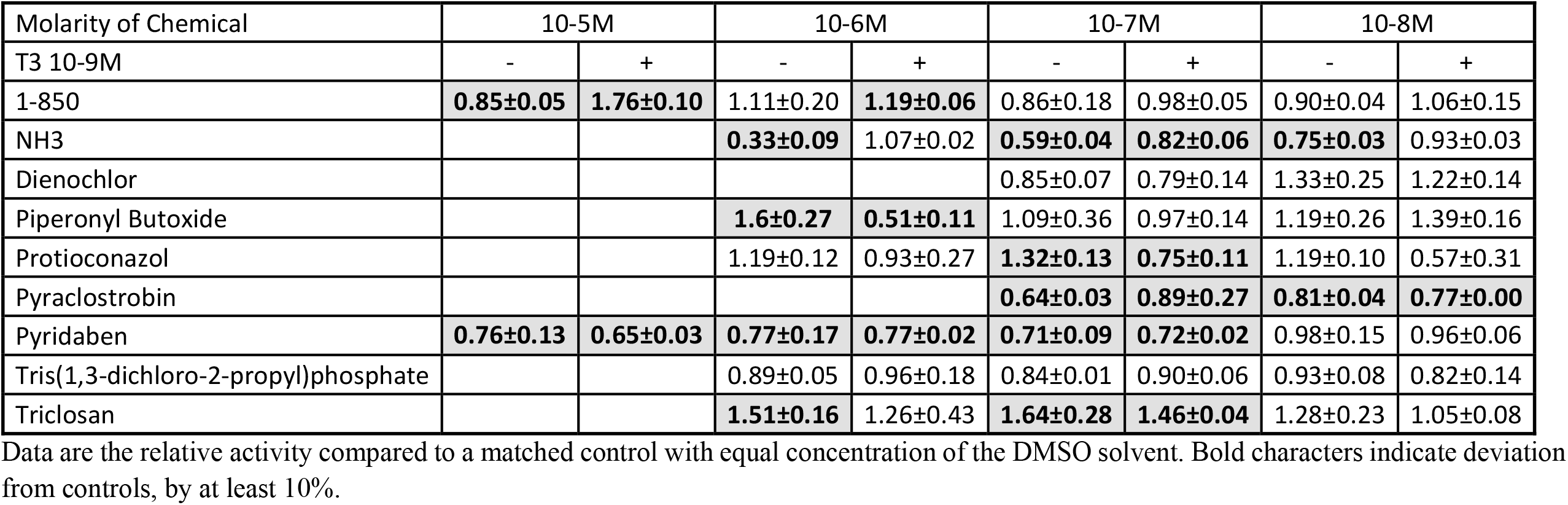
Capacity to displace the NcoR corepressor from the TRα1 ligand binding domain.

| Molarity of Chemical | 10-5M |  | 10-6M |  | 10-7M |  | 10-8M |  |
| --- | --- | --- | --- | --- | --- | --- | --- | --- |
| T3 10-9M | - | + | - | + | - | + | - | + |
| 1-850 | **0.85±0.05** | **1.76±0.10** | 1.11±0.20 | **1.19±0.06** | 0.86±0.18 | 0.98±0.05 | 0.90±0.04 | 1.06±0.15 |
| NH3 |  |  | **0.33±0.09** | 1.07±0.02 | **0.59±0.04** | **0.82±0.06** | **0.75±0.03** | 0.93±0.03 |
| Dienochlor |  |  |  |  | 0.85±0.07 | 0.79±0.14 | 1.33±0.25 | 1.22±0.14 |
| Piperonyl Butoxide |  |  | **1.6±0.27** | **0.51±0.11** | 1.09±0.36 | 0.97±0.14 | 1.19±0.26 | 1.39±0.16 |
| Protioconazol |  |  | 1.19±0.12 | 0.93±0.27 | **1.32±0.13** | **0.75±0.11** | 1.19±0.10 | 0.57±0.31 |
| Pyraclostrobin |  |  |  |  | **0.64±0.03** | **0.89±0.27** | **0.81±0.04** | **0.77±0.00** |
| Pyridaben | **0.76±0.13** | **0.65±0.03** | **0.77±0.17** | **0.77±0.02** | **0.71±0.09** | **0.72±0.02** | 0.98±0.15 | 0.96±0.06 |
| Tris(1,3-dichloro-2-propyl)phosphate |  |  | 0.89±0.05 | 0.96±0.18 | 0.84±0.01 | 0.90±0.06 | 0.93±0.08 | 0.82±0.14 |
| Triclosan |  |  | **1.51±0.16** | 1.26±0.43 | **1.64±0.28** | **1.46±0.04** | 1.28±0.23 | 1.05±0.08 |

Data are the relative activity compared to a matched control with equal concentration of the DMSO solvent. Bold characters indicate deviation from controls, by at least 10%.

**Table 4.** RT-qPCR measurement of *Hairless* and *Klf9* mRNA T3 response

| Molarity of Chemical |  | 10-5M | 10-6M | 10-7M | 10-8M | 10-9M |
| --- | --- | --- | --- | --- | --- | --- |
| 1-850 | *Hairless* | **0.45** |  |  |  |  |
|  | *Klf9* | **0.57** |  |  |  |  |
| NH3 | *Hairless* |  | **0.58** |  |  |  |
|  | *Klf9* |  | **0.58** |  |  |  |
| Azoxystrobin | *Hairless* |  | 0.93 | 0.92 |  |  |
|  | *Klf9* |  | 1.25 | 1.38 | 0.61 |  |
| Dienochlor | *Hairless* |  | 0.87 | 0.84 | 0.66 |  |
|  | *Klf9* |  | **0.41** | **0.45** | 0.97 |  |
| Fenitrothion | *Hairless* |  | 1.10 | 1.18 | 1.97 |  |
|  | *Klf9* |  | 0.94 | 1.18 | 1.66 |  |
| Phosalone | *Hairless* |  | 1.26 | 1.66 |  |  |
|  | *Klf9* |  | 1.20 | 1.25 |  |  |
| Picoxystrobin | *Hairless* |  | 2.68 | 1.17 | 1.69 |  |
|  | *Klf9* |  | 1.20 | 1.35 | 1.33 |  |
| Piperonyl Butoxide | *Hairless* |  | 1.10 | 0.93 | 1.04 |  |
|  | *Klf9* |  | 1.07 | 1.00 | 1.31 |  |
| Pyraclostrobin | *Hairless* |  | 1.26 | 2.42 | 1.78 |  |
|  | *Klf9* |  | 0.96 | 1.18 | 1.19 |  |
| Quinoxifen | *Hairless* |  | 0.93 | 0.99 |  |  |
|  | *Klf9* |  | 0.89 | 0.96 |  |  |
| Trifloxystrobin | *Hairless* |  | 1.19 | 1.04 | 0.82 | 0.36 |
|  | *Klf9* |  | 1.19 | 1.16 | 1.23 | 1.14 |

Relative response of mRNA levels compared to control without pesticide.

### Transcriptome analysis

To gain a broader and unbiased view of the influence of the complex response of neural cells to the tested chemicals, we selected the 7 pesticides which displayed the most visible effects in the reporter tests for a genome wide analysis of gene expression: pyperonyl butoxide, pyridaben, emanectin benzoate, and 3 strobilurins (picoxystrobin, pyraclostrobin and trifloxystrobin). We first used human neuroblastoma cell line SH-SY5Y, and whole genome Ampliseq to analyze the response to the pesticides, to T3, and the interference between the two responses. Except for piperonyl butoxide, all compounds had a significant influence on gene expression in these cells (Figure 1A). However, there was no indication that TH response was altered as most of the disrupted genes are not responsive to T3. The only exception was for pyraclostrobin, which exerts a broad influence on gene expression including an interference with the TH response for some of the T3 responsive genes (Figure 1B). However SH-SY5Y only display moderate TH response, they might not be well suited to identify a minor alteration of this response. We thus repeated the experiment for pyraclostrobin and piperonyl butoxide on mouse C17.2α neural cells, as these compounds were active in all reporter tests, but displayed a puzzling pattern. RNA-seq confirmed that these cells, which have been engineered to overexpress TRα1, displayed a robust TH response (Guyot *et al.* 2014, Chatonnet *et al.* 2013). Piperonyl butoxide had a detectable influence on gene expression. Although T3 had a moderate effect on the response of genes to piperonyl butoxide (Figure 2A) there was no visible reciprocal influence of piperonyl butoxide on T3 response (Figure 2A,B). Pyraclostrobin alters the expression of a larger set of genes (Figure 2C). Although this pesticide does not systematically alters the cellular response to T3, it potentiates the action of T3 for a subset of T3 responsive genes, and exert the opposite effect for another subset of these genes (Figure 2D) as already supported by the previous analysis on SH-SY5Y cells. The significance of the influence of the two pesticides on T3 response was addressed by using Geneset enrichment analysis (GSEA). We used here this sensitive non-parametric method to analyse the distribution of the T3 responsive genes within the list of all expressed genes, ranked according to their response to pesticide. The list of genes up-regulated by T3 in the present experiment was first defined by differential expression analysis (Deseq2 2 factors T3 and pesticide). GSEA showed no significant effect of piperonyl butoxide on these genes in absence of T3. It also indicated a statistically significant influence of pyraclostrobin on T3 responsive genes (p-value: 0.02) with a trend toward an agonist-like effect (Suppl. figure S2A). As already observed in the clustering analysis, this effect only concerns a subset of the T3 responsive genes.

**Figure 1:**
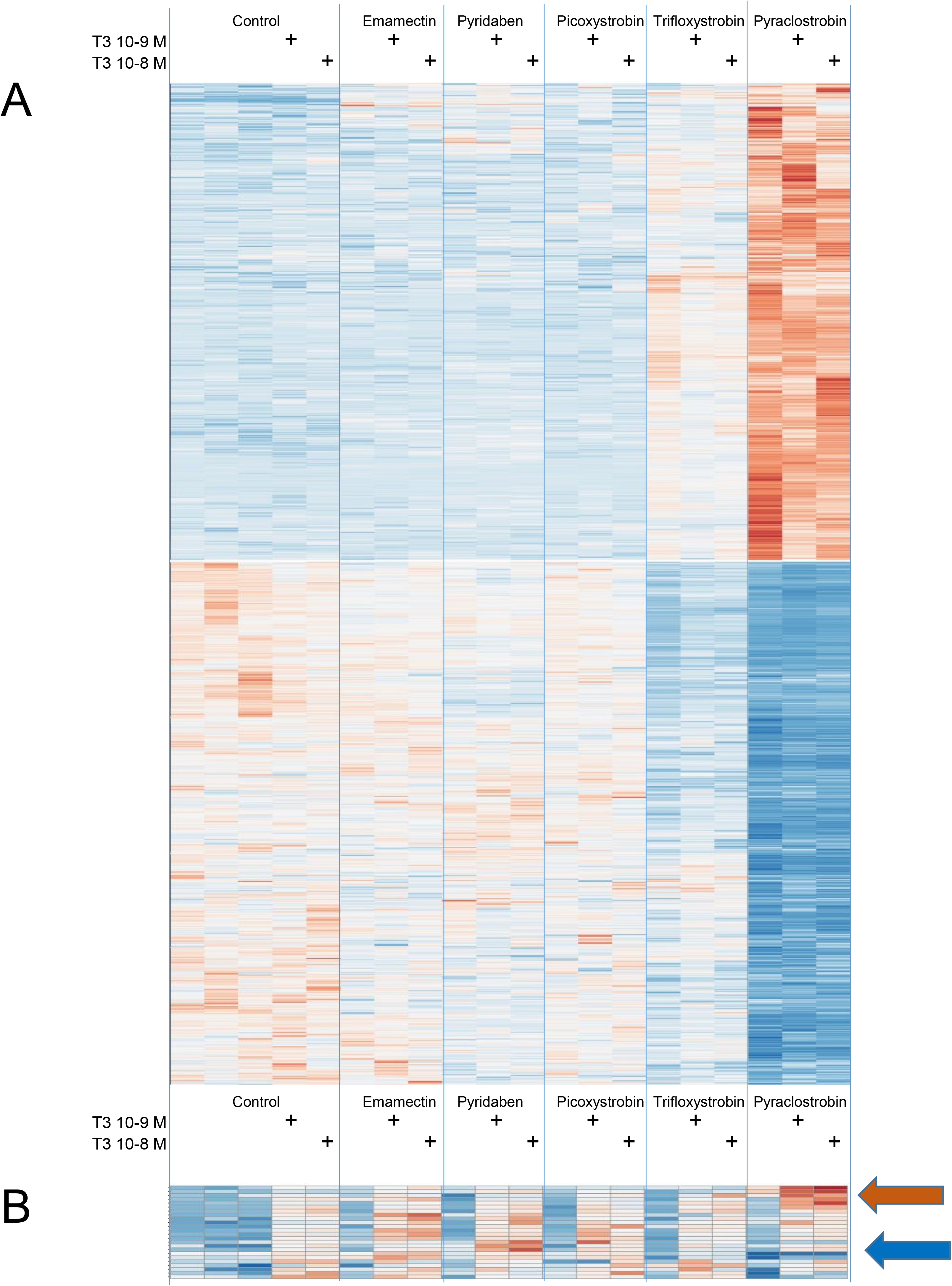
Genome wide analysis of the pesticides and thyroid hormone response of SHSY5Y human neuroblastoma cells. The heatmaps represent the result of a clustering analysis for the response of SHSY5Y human neuroblastoma cells to pesticides and thyroid hormone (red up-regulation, blue down-regulation). Ampliseq results obtained from cells exposed to a pesticide (10^−6^M), in absence or in presence of T3 at the indicated concentration were submitted to differential expression analysis (Deseq2; First factor pesticide, second factor: T3. Adjusted p-value <0.05) identified a number of genes which expression is sensitive to the presence of pesticide (Emamectin: 336; Pyridaben: 225; Picoxystrobin 57; Trifloxystrobin: 5856; Pyraclostrobin: 7290). The reciprocal analysis identified 25 T3 responsive genes. A. Pesticide response. Only the 1130 genes displaying at least a 2-fold response to one of the pesticides are included. B. T3 response. No systematic bias is observed, which would be expected if one of the pesticide was a TRα1 ligand. Note however that the response to T3 of a subset of genes is sensitive to pyraclostrobin. A subset of genes becomes more sensitive to T3 stimulation (red arrow), while the T3 response of others is dampened (blue arrow).

**Figure 2:**
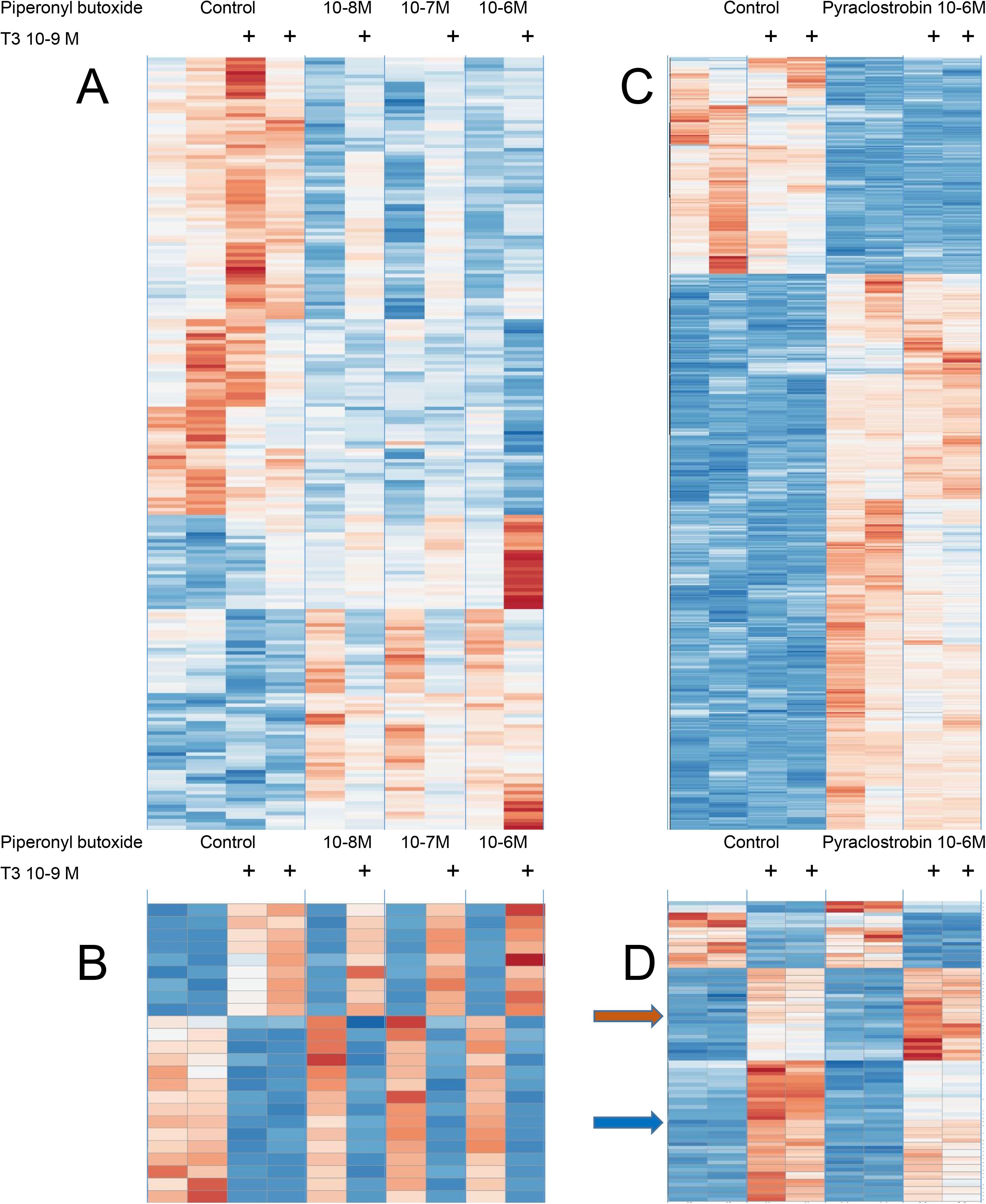
Transcriptome response of C17.2α cells to piperonyl butoxide, pyraclostrobin and thyroid hormone. A-B) Ampliseq results obtained from cells exposed to piperonyl butoxide at different concentrations, in absence of in presence of T3. Differential expression analysis (Deseq2; First factor pesticide, second factor: T3. Adjusted p-value <0.05) identified a number of genes which expression is sensitive to the presence of piperonyl butoxide (10^−8^ M n=51; 10^−7^ M n=75; 10^−6^ M n=197). The reciprocal analysis identified 25 T3 responsive genes. A) Pesticide response. B) T3 response. The clustering analysis does not highlight a systematic bias in T3 response, which would be expected if one of the pesticide was a TRα1 ligand. Note however that the response to piperonyl butoxide is clearly different in present of T3. C-D) RNA-seq results obtained from cells exposed to Pyraclostrobin (10^−6^ M) in absence of in presence of T3. (red up-regulation, blue down-regulation). Only genes with at least a 2 fold-change in expression for one pesticide concentration are plotted (630 out of 4780). Note that the sensitivity to T3 is increased for a subset of genes (red arrow) and decreased for another subset (blue arrow).

In all the previous experiments, pyraclostrobin stands out as a more active strobilurin. Whether the other strobilurins would exert a similar effect at higher concentrations, or are acting on different pathways, remains unclear. To address this question, we performed another RNA-seq analysis, testing the influence of the four strobilurines at different concentrations in the absence of T3. This analysis showed that the responses to the different strobilurins do not fully overlap. In particular, picoxystrobin is also active at low concentration, changing the expression of a distinct subset of genes (Figure 3).

**Figure 3:**
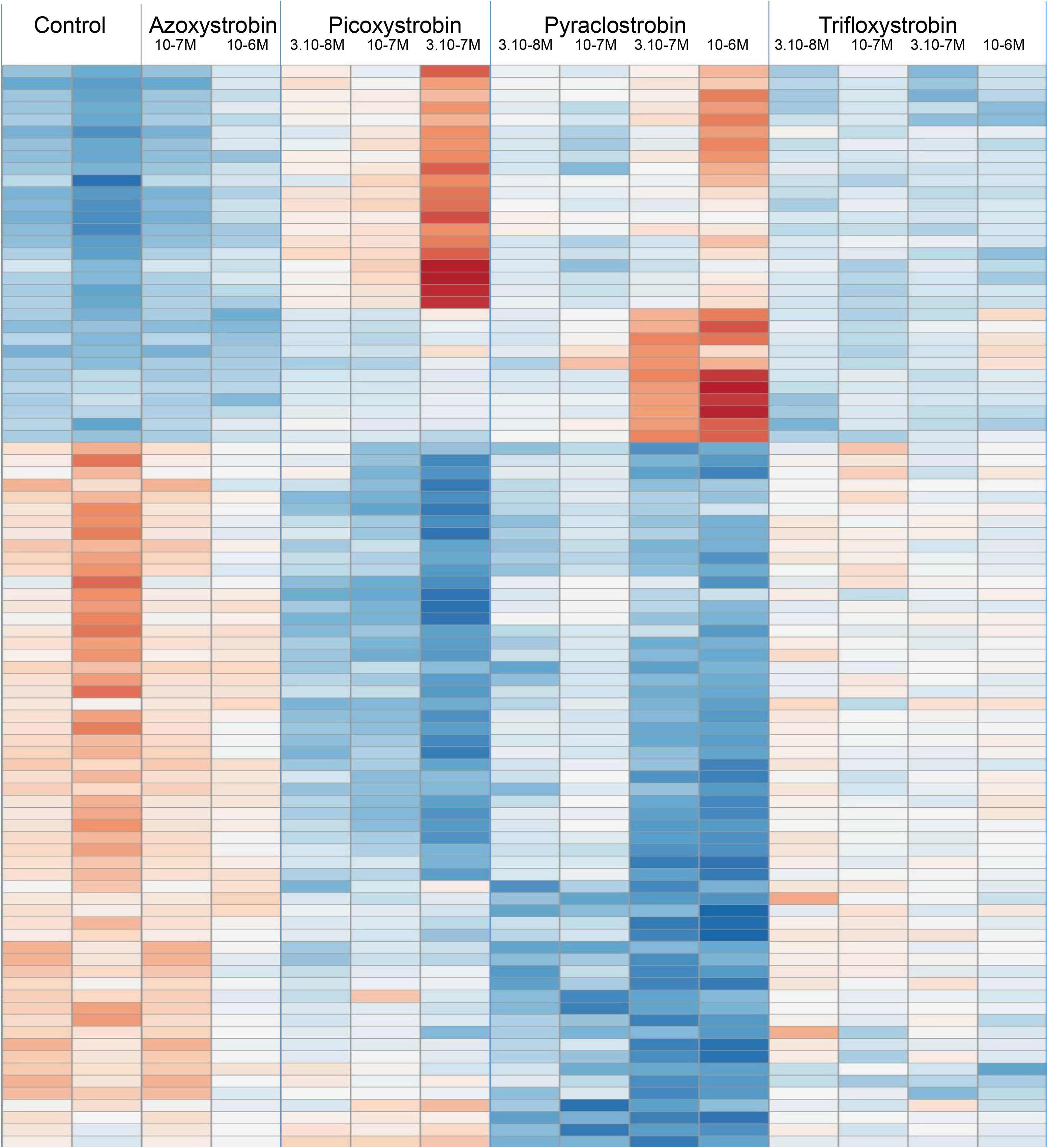
Transcriptome response of C17.2α cells to strobilurins in absence of thyroid hormone. Differentially expressed genes were identified by comparing all exposed samples to controls (Deseq2 1 factor, adjusted p-value <0.05) after exposing C17.2α to growing concentrations of either azoxystrobin (n=4), picoxystrobin(n=3065), pyraclostrobin (n=962), or trifloxystrobin (n=272). Only the 90 genes for which a fold-change >2 was measured for at least one pesticide were included in the clustering analysis (red: up-regulation, blue: down-regulation).

### Mining published data for putative TH disruptors

The above data converge to suggest that none of the tested chemicals is a genuine agonist or antagonist of TRα1. However, some molecules, notably pyraclostrobin, interfere with T3 signaling in an indirect manner, modifying the response of a specific subset of TRα1 target genes. The above analysis also shows that GSEA can identify such molecules in an unbiased manner, even if the T3 response is only partially altered. We thus took advantage of the availability of a large set of RNA-seq data (GSE70249) which analyzed the influence of 297 chemicals, mainly pesticides, on primary cell cultures prepared from the cortex of newborn mice (Pearson *et al.* 2016). Because the culture medium used in these experiments contained T3, present in the B27 supplement and serum (approximatively 10^−9^M), we believe that it is mainly suitable to detect TRα1 antagonists, although agonists might potentiate the effect of T3. The response to T3 of this cellular model has been fully characterized by RNA-seq in a different study (Gil-Ibanez *et al.* 2015), allowing us to define the top 100 T3 induced genes in this system, all of them having a minimal fold-change above 1.5 (pval <0.01, FDR < 0.05). The normalized abundance matrix was used to rank 13304 genes from the most up-regulated to the most down-regulated in response to each compound. This analysis converged to identify some of the compounds that we tested in reporter cells as active on T3 signaling, with antagonist-like properties (emamectin benzoate, piperonyl butoxide, propiconazole and Pyraclostrobin (Table S3). Clustering analysis of the T3 responsive genes for the 33 compounds that we tested in reporter assays showed however that pyraclostrobin and trifloxystrobin are the only chemicals clearly segregating from controls (Figure 4). Again, when considering the full dataset, only a small group of chemical including pyraclostrobin and trifloxystrobin branched out of the controls (Suppl. figure S4). Based on this GSEA analysis (Suppl. Figure S2B), we selected 6 compounds, predicted to significantly interfere with TH signaling. Although our *in vitro* tests revealed that these compounds are active at different concentrations, they do not suggest that any of these is a genuine TRα1 agonist or antagonist (Table S5).

**Figure 4:**
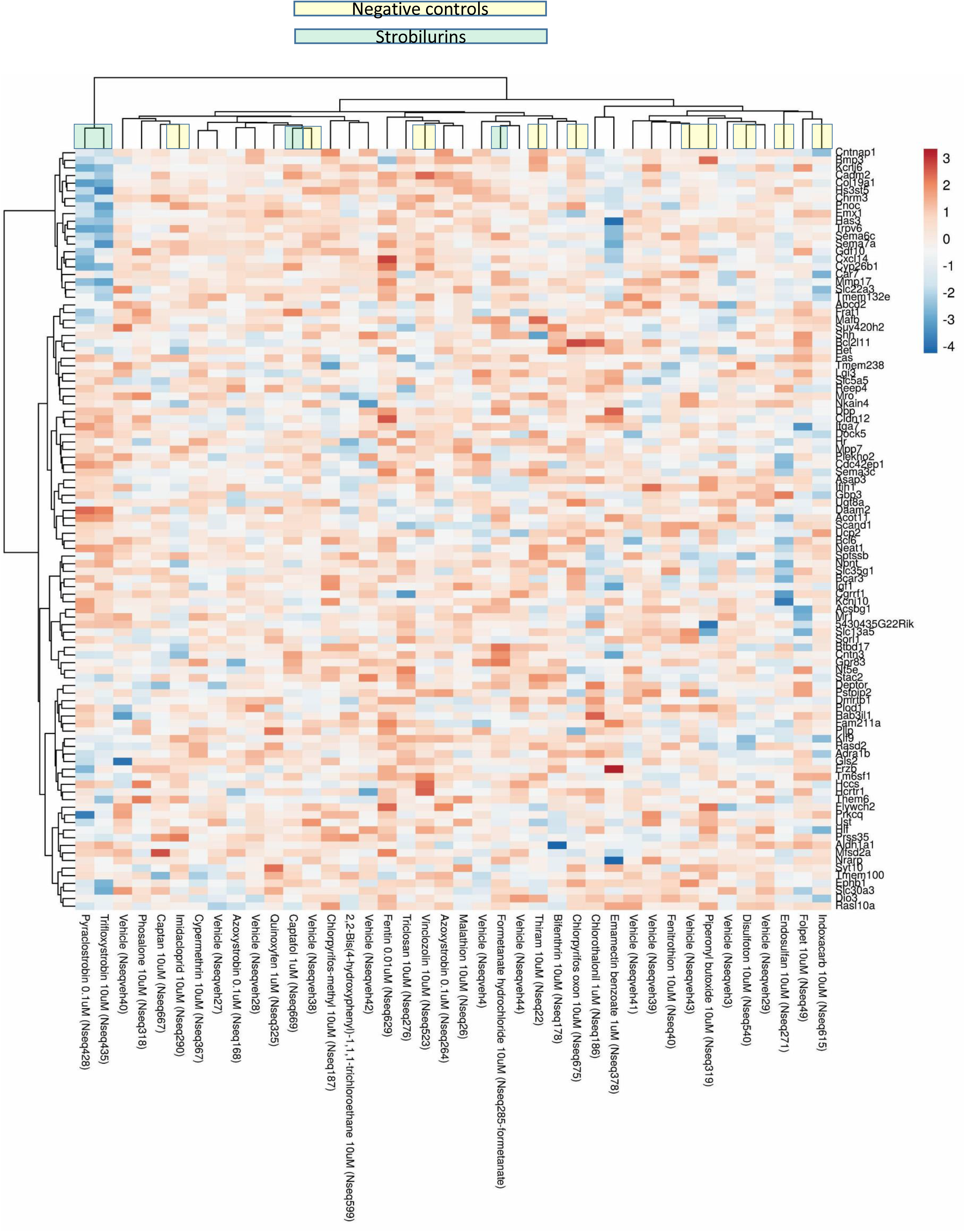
Pesticides activity on cortical neurons. A 2D-clustering analysis was performed with the normalized expression data extracted from published results (Pearson *et al.* 2016), which studied the influence of pesticides on primary cultures of mouse cortical neurons. We extracted the data for the 33 pesticides tested in the present study, and 10 negatives controls (yellow boxes). Not that pyraclostrobin and trifloxystrobin are the only strobilurins (blue boxes) to branch out of the controls on the upper dendrogram.

### T3 independent action of strobilurins

The transcriptome analyses that we performed highlight the capacity of strobilurins to exert a major influence on gene expression in neural cells, which is unrelated to T3 signaling. Gene Ontology analysis (http://cbl-gorilla.cs.technion.ac.il/) notably indicates that high concentrations of pyraclostrobin down-regulates the expression of genes encoding major component of nucleosome assembly, notably histones. As we suspected that this would translate in an inhibition of cell proliferation, we tested the capacity of strobilurins to influence C17.2α cells growth (Suppl. figure S6). Interestingly, this indicate that the four tested strobilurins tend to favor cell growth at low concentration. By contrast high concentrations of pyraclostrobin, and to a lower extend trifloxystrobin, had the opposite effect. These effects were not sensitive to the presence of T3. Taken together, these observations suggest that one major outcome of the broad perturbation of gene expression selectively induced by by pyraclostrobin and trifloxystrobin is a reduction in cell growth.

## Discussion

We present here an extensive assessment of the capacity of pesticides to interfere with the cellular response to T3. One main conclusion is that none of the tested compound is behaving as a genuine TRα1 agonist or antagonist. This reinforces the conclusion of the Tox21 screen, performed on 8305 compounds, according to which many environmental chemicals do not act as TRα1 ligand at non-toxic concentrations, either as agonists or antagonists (Paul-Friedman *et al.* 2019). This contrasts with other nuclear receptors, for which a number of environmental ligands have been identified (Toporova *et al.* 2020, Liu *et al.* 2019). This peculiarity probably reflects the small size of the ligand binding pocket of TRs, and the specific chemical properties of their natural iodinated ligands, T4 and T3. However the presence of halogens in the chemical structure (fluoride, iodine, chloride, or bromide) is not an indication for a possible binding to TRs, as halogens are frequent is pesticides, and absent from the NH-3 synthetic TRα1 ligand. This study also illustrates the importance of combining several *in vitro* assays to draw firm conclusions on the mode of action of THD, as previously recommended by others (Paul-Friedman *et al.* 2019), as single endpoints screens produce a high rate of false positives.

Our data illustrate the benefit provided by transcriptome analysis for *in vitro* toxicology: first it is unbiased, and able to capture any unexpected alteration in the cellular physiology. Second, if performed on relevant cellular models, it helps to prioritize some chemicals for *in vivo* assessment, and limit the use of animals, as recommended by ethical guidelines. Third, as the expression level of each gene of a given signaling pathway can be considered as an independent estimate of the signaling level, it provides outstanding statistical power to detect minor effects on defined pathways. While the tested pesticides have only a marginal influence of TH signaling, our genome wide analysis shows that many have the potential to compromise neurodevelopment, by exerting a broad influence on gene expression in neural cells, even at low concentration. In particular, strobilurins, and notably pyraclostrobin, stand out as the most active compounds that we have tested, in agreement with previous conclusions (Pearson *et al.* 2016). These chemicals were designed to act as fungicides, selectively inhibiting the mitochondrial cytochrome-bc1 complex of fungi. They are however toxic on zebrafish embryo at low concentration (LC50 1.5×10^−7^M for pyraclostrobin) (Zhang *et al.* 2017). In mice, a significant reduction in body weight is also observed upon chronic exposure if the food contains more than 10 mg/kg of pyraclostrobin (Bartholomaeus 2003). The neural cells response to pyraclostrobin and trifloxystrobin expands to a fraction of the T3 responsive genes, but is mainly T3 independent, and disturb the cell growth. Whether these pesticides can reach the foetal brain during pregnancy, which is currently unknown.

More generally, our study suggests that the definition of the second class of THD, which alter the cellular response to T3, should be reconsidered. Some chemicals do not directly interfere with TRs function, but have the capacity to modify the response to T3 for a fraction of the TRs target genes only. If these genes are key mediators of the neurodevelopmental function of T3, this should be a matter of concern.

### Limitations of the study

In vitro analysis is well suited for mechanistic analysis, but insufficient to predict the outcome of in vivo toxicity. In particular, it does not take into account the generation of secondary metabolites after catabolism and does not inform on the distribution of xenobiotics in the organisms.

## Acknowledgements

We thank Chloé Morin, Nathan Gay, Manon Baudry, Julie Renaud and Juna Cuomo for help in luciferase and qRT-PCR assays. We thank Benjamin Gillet and Sandrine Hughes from the IGFL-PSI facility for deep sequencing. This work was supported by ANSES (Thyrogenox program, PNRPE Ecophyto II) and the European Union’s Horizon 2020 research and innovation program under grant agreement no. 825753 (ERGO).

## Supplementary data

## Legends of supplementary figures

**Figure S1:**
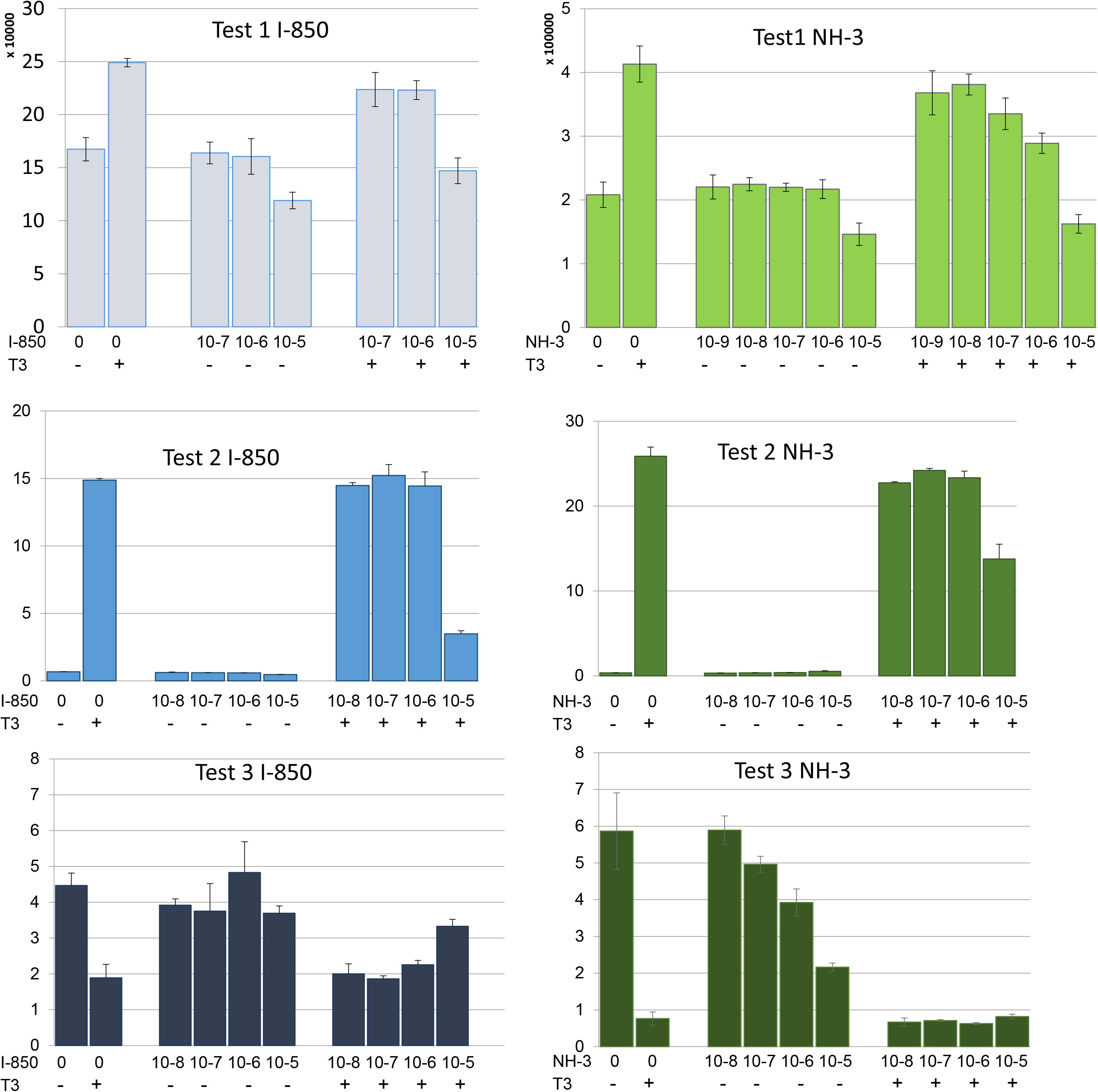
Validation of reporter tests with reference compounds. T3 increases the luciferase activity of C17.2α-Hrluc cells (upper panel) and HEK293–Gal4TRα1luc cell line (middle panel). 1-850 and NH-3 are TRα1 synthetic ligands which inhibit T3 mediated transactivation in a concentration dependent manner. In HEK293 cells co-transfected with pBKGal4NcoR/pBKVP16TRα1/pGal4REx5-βglob-luc-SVNeo the interaction between the NcoR corepressor domain and the TRα1 ligand binding domain in absence of T3 results in a transactivation of the UAS driven promoter and a high luciferase activity. Addition of T3 destabilizes the interaction between the two hybrid proteins and reduces luciferase activity. 1-850 has little effect in absence of T3, and prevents T3 response in a dose dependent manner. NH-3 per se destabilizes the interaction, and cooperate with T3 to reduce luciferase activity.

**Figure S2:**
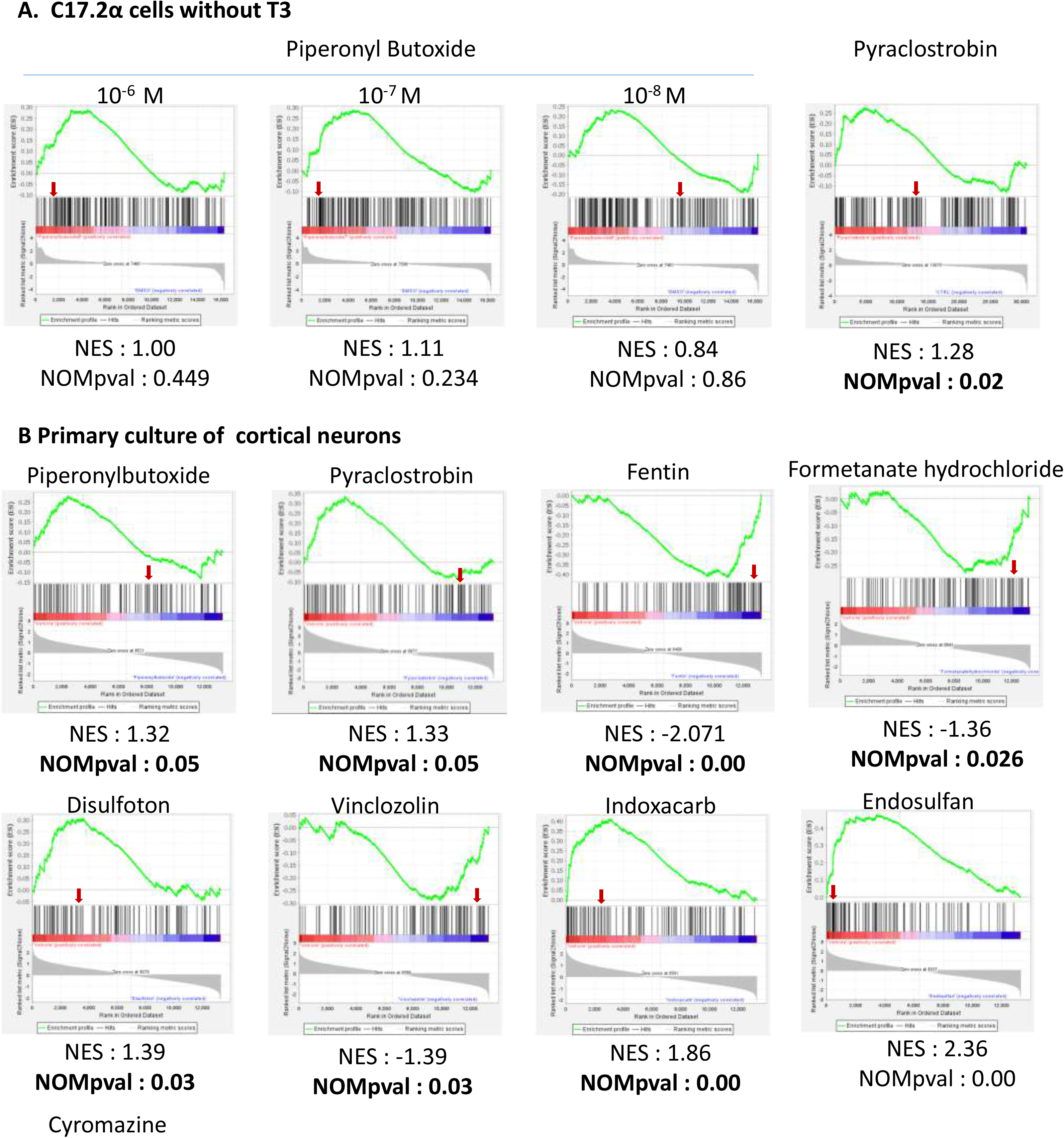
Geneset enrichment analysis. The geneset is the group of T3 responsive genes, as defined by differential analysis on C17.2α (A) or on primary cultures of cortical neurons (B). Vertical black bars represent the position of each T3 responsive gene in the list of genes ranked by expression level. The green curve represents The distribution is significantly shifted toward high expression (adjusted p-value <0.05) indicating the predominance of an agonist-like activity of Pyraclostrobin. The green curve corresponds to the ES (enrichment score) curve, which is the running sum of the weighted enrichment score obtained from GSEA software, while the normalized enrichment score (NES) and the corresponding P-value are reported within each graph. The red arrow indicates the position of Hairless, a gene which expression is highly sensitive to T3.

**Figure S3:**
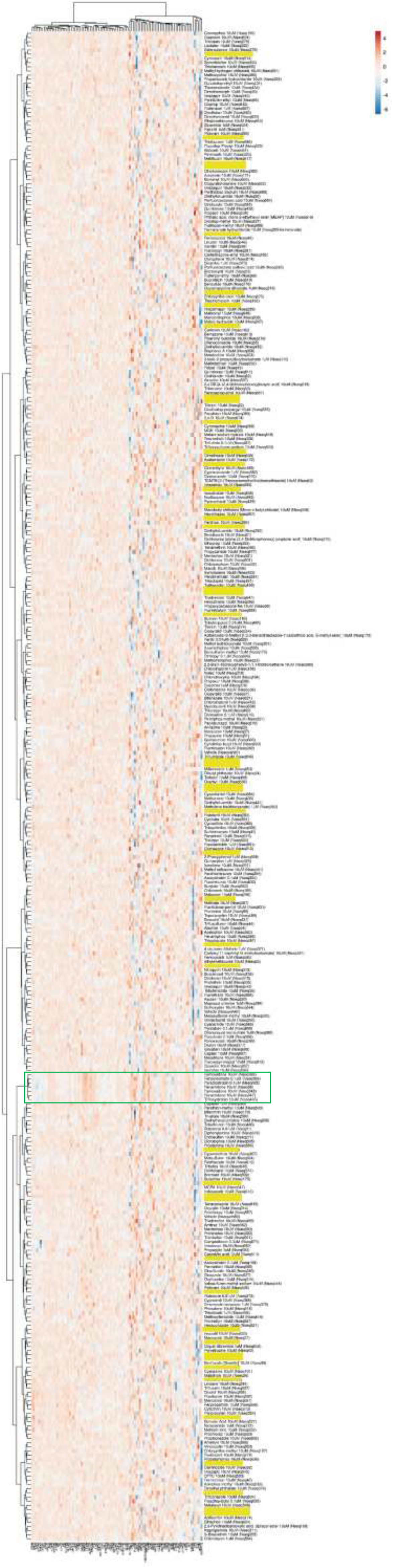
2D Clustering of the RNA-seq data of T3 responsive genes for all pesticides tested on primary cultures of cortical neurons. The yellow boxes correspond to negative controls. The green frame highlights a cluster of 5 pesticides which alter the expression of a small group of T3 responsive gene. This cluster contains an acaricide (fenpyroximate) and 4 fungicides (Pyraclostrobin, trifloxystrobin, famoxadone and fenamidone) which all inhibit the mitochondria electron transport chain of their target organisms.

**Figure S6:**
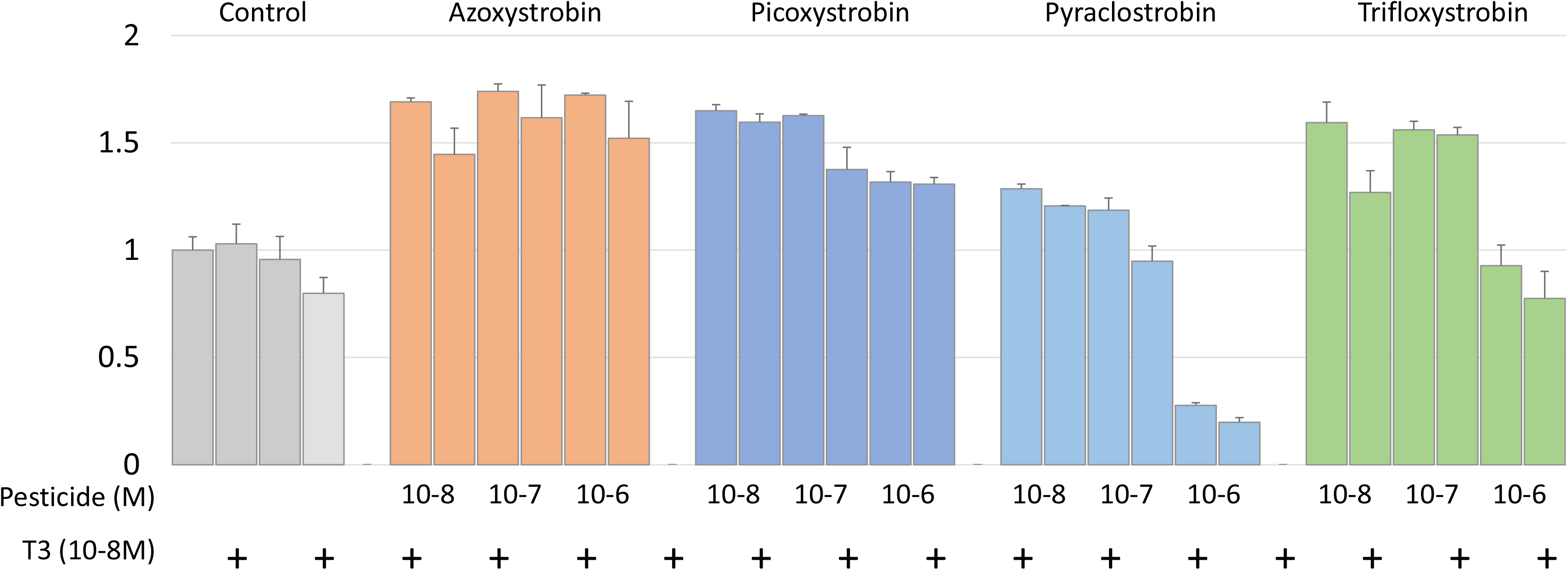
C17.2α cell growth in presence of strobilurins. Cells were seeded in 96 wells (5000 cells/well) in medium prepared with hormone-depleted serum and supplemented with the indicated molarity of pesticide and T3. Cell growth was quantified 4 days later with CellTiter-Glo Luminescent Cell Viability Assay. While all strobilurin tend to favor cell growth at low concentration, pyraclostrobine and trifloxystrobine have the opposite effect when used at high concentration.

**Table S3.** GSEA analysis of selected pesticides properties on primary cultures of cortical neurons.

|  | Rank of genes up-regulated by T3 |  |  |  | Expected activity |
| --- | --- | --- | --- | --- | --- |
| Compound | Enrichment Score | Normalized<br>Enrichment Score | Nom. p-value | rank at max |  |
| Azoxystrobin | 0.21 | 0.95 | 0.54 | 2848 |  |
| Bifenthrin | -0.23 | -1.14 | 0.20 | 1746 |  |
| Captafol | -0.22 | -1.08 | 0.29 | 3185 |  |
| Captan | 0.19 | 0.91 | 0.65 | 426 |  |
| Chlorothalonil | 0.27 | 1.21 | 0.12 | 3035 |  |
| Chlorpyrifos | -0.23 | -1.12 | 0.22 | 3226 |  |
| Cypermethrin | -0.22 | -1.11 | 0.23 | 3430 |  |
| Emamectinbenzoate | 0.32 | 1.37 | 0.05 | 1793 | antagonist |
| Fenitrothion | 0.21 | 0.95 | 0.55 | 2424 |  |
| Folpet | 0.23 | 1.08 | 0.28 | 2294 |  |
| Imidacloprid | 0.24 | 1.16 | 0.17 | 2398 |  |
| Malathion | -0.17 | -0.86 | 0.77 | 3582 |  |
| Phosalone | 0.23 | 1.08 | 0.29 | 2636 |  |
| Piperonylbutoxide | 0.28 | 1.32 | 0.05 | 2427 | antagonist |
| Propiconazole | 0.32 | 1.54 | 0.01 | 2353 | antagonist |
| Pyraclostrobin | 0.33 | 1.34 | 0.05 | 2868 | antagonist |
| Pyridaben | 0.25 | 0.94 | 0.58 | 2356 |  |
| Quinoxifen | -0.23 | -1.14 | 0.21 | 4264 |  |
| Thiram | -0.21 | -0.90 | 0.69 | 2457 |  |
| Triclosan | -0.23 | -1.15 | 0.18 | 2259 |  |
| Trifloxystrobin | 0.32 | 1.29 | 0.08 | 3125 |  |
| Disulfoton | 0.31 | 1.39 | 0.04 | 3571 | antagonist |
| Endosulfan | 0.48 | 2.36 | 0.00 | 3434 | antagonist |
| Fentin | -0.42 | -2.07 | 0.00 | 2387 | agonist |
| Formetanatehydrochloride | -0.28 | -1.36 | 0.03 | 4507 | agonist |
| Indoxacarb | 0.41 | 1.86 | 0.00 | 2978 | antagonist |
| Vinclozolin | -0.29 | -1.39 | 0.03 | 3395 | agonist |

**Table S4.** GSEA analysis of selected pesticides properties on primary cultures of cortical neurons.

| Compound | Rank of genes up-regulated by T3 |  |  |  | Expected activity |
| --- | --- | --- | --- | --- | --- |
|  | Enrichment Score | Normalized | Nom. p-value | rank at max |  |
| Azoxystrobin | 0.21 | 0.95 | 0.54 | 2848 |  |
| Bifenthrin | -0.23 | -1.14 | 0.20 | 1746 |  |
| Captafol | -0.22 | -1.08 | 0.29 | 3185 |  |
| Captan | 0.19 | 0.91 | 0.65 | 426 |  |
| Chlorothalonil | 0.27 | 1.21 | 0.12 | 3035 |  |
| Chlorpyrifos | -0.23 | -1.12 | 0.22 | 3226 |  |
| Cypermethrin | -0.22 | -1.11 | 0.23 | 3430 |  |
| Emamectinbenzoate | 0.32 | 1.37 | **0.05** | 1793 | antagonist |
| Fenitrothion | 0.21 | 0.95 | 0.55 | 2424 |  |
| Folpet | 0.23 | 1.08 | 0.28 | 2294 |  |
| Imidacloprid | 0.24 | 1.16 | 0.17 | 2398 |  |
| Malathion | -0.17 | -0.86 | 0.77 | 3582 |  |
| Phosalone | 0.23 | 1.08 | 0.29 | 2636 |  |
| Piperonylbutoxide | 0.28 | 1.32 | **0.05** | 2427 | antagonist |
| Propiconazole | 0.32 | 1.54 | **0.01** | 2353 | antagonist |
| Pyraclostrobin | 0.33 | 1.34 | **0.05** | 2868 | antagonist |
| Pyridaben | 0.25 | 0.94 | 0.58 | 2356 |  |
| Quinoxifen | -0.23 | -1.14 | 0.21 | 4264 |  |
| Thiram | -0.21 | -0.90 | 0.69 | 2457 |  |
| Triclosan | -0.23 | -1.15 | 0.18 | 2259 |  |
| Trifloxystrobin | 0.32 | 1.29 | 0.08 | 3125 |  |
| Disulfoton | 0.31 | 1.39 | **0.04** | 3571 | antagonist |
| Endosulfan | 0.48 | 2.36 | **0.00** | 3434 | antagonist |
| Fentin | -0.42 | -2.07 | **0.00** | 2387 | agonist |
| Formetanatehydrochloride | -0.28 | -1.36 | **0.03** | 4507 | agonist |
| Indoxacarb | 0.41 | 1.86 | **0.00** | 2978 | antagonist |

**Table S5:** Testing 6 compounds predicted to be active on T3 signaling by GSEA.

|  |  |  |  |  |  |  |  |  |
| --- | --- | --- | --- | --- | --- | --- | --- | --- |
| Molarity of Chemical | 10-5M |  | 10-6M |  | 10-7M |  | 10-8M |  |
| T3 10-9M | - | + | - | + | - | + | - | + |
| Predicted agonists |  |  |  |  |  |  |  |  |
| Test 1 |  |  |  |  |  |  |  |  |
| Fentin hydroxide |  |  |  |  | 1.01±0.18 | 0.68±0.05 | 0.94±0.11 | 0.78±0.02 |
| Formetanate Hydrochloride | 1.14±0.15 | 0.87±0.08 | 1.13±0.07 | 0.89±0.08 | 1.06±0.07 | 0.96±0.09 | 1.01±0.05 | 1.01±0.16 |
| Vinclozolin | 0.96±0.07 | 0.71±0.02 | 1.00±0.24 | 0.76±0.02 | 1.01±0.02 | 0.76±0.06 | 1.11±0.11 | 0.56±0.02 |
| Test 2 |  |  |  |  |  |  |  |  |
| Fentin hydroxide |  |  |  |  | 2.25±0.14 | 0.24±0.01 | 1.86±0.18 | 0.39±0.05 |
| Formetanate Hydrochloride | 1.17±0.08 | 1.38±0.05 | 1.04±0.09 | 1.14±0.07 | 0.97±0.02 | 1.23±0.03 | 0.97±0.04 | 1.26±0.05 |
| Vinclozolin | 0.96±0.07 | 0.71±0.02 | 1.00±0.24 | 0.76±0.02 | 1.01±0.02 | 0.76±0.06 | 1.11±0.11 | 0.56±0.02 |
| Test 3 |  |  |  |  |  |  |  |  |
| Fentin hydroxide |  |  |  |  | 0.17±0.01 | 0.16±0.01 | 0.51±0.07 | 0.52±0.04 |
| Formetanate Hydrochloride | 0.88±11 | 0.71±0.10 | 0.98±0.18 | 0.70±0.02 | 0.93±0.08 | 0.84±0.04 | 1.05±0.11 | 0.93±0.02 |
| Vinclozolin |  |  |  |  |  |  |  |  |
| Predicted antagonists |  |  |  |  |  |  |  |  |
| Test 1 |  |  |  |  |  |  |  |  |
| Disulfoton | 0.72±0.07 | 0.72±0.05 | 0.92±0.13 | 0.84±0.09 | 0.92±0.07 | 0.77±0.05 | 0.97±0.04 | 0.83±0.07 |
| Beta Endosulfan |  |  | 1.01±0.13 | 0.78±0.07 | 1.15±0.19 | 0.99±0.10 | 1.02±0.05 | 0.97±0.07 |
| Indoxacarb |  |  | 0.99±0.14 | 0.92±0.10 | 1.12±0.10 | 0.91±0.10 | 1.04±0.15 | 0.85±0.04 |
| Test 2 |  |  |  |  |  |  |  |  |
| Disulfoton | 1.56±0.10 | 0.84±0.12 | 0.67±0.07 | 0.63±0.01 | 0.85±0.45 | 0.74±0.08 | 1.60±0.01 | 0.70±0.05 |
| Beta Endosulfan | 0.88±0.09 | 0.99±0.04 | 0.92±0.07 | 0.93±0.08 | 0.90±0.01 | 0.97±0.05 | 0.97±0.05 | 1.01±0.05 |
| Indoxacarb |  |  | 1.04±0.02 | 0.91±0.03 | 0.98±0.07 | 0.90±0.05 | 1.02±0.11 | 0.90±0.09 |
| Test 3 |  |  |  |  |  |  |  |  |
| Disulfoton |  |  |  |  |  |  |  |  |
| Beta Endosulfan |  |  | 1.19±0.15 | 1.20±0.07 | 1.11±0.04 | 1.22±0.26 | 1.04±0.04 | 1.15±0.07 |
| Indoxacarb |  |  | 0.90±0.08 | 0.89±0.05 | 1.09±0.08 | 1.11±0.08 | 1.07±0.11 | 1.17±0.07 |

## Notes

### Competing Interest Statement

The authors have declared no competing interest.

